# Baicalin Ameliorates Defective Decidualization in URSA by Regulating Mitochondrial Fission Induced Necroptosis

**DOI:** 10.1101/2023.03.29.534851

**Authors:** Xiaoxuan Zhao, Ying Zhao, Qujia Yang, Jing Ma, Yang Zhao, Suxia Wang, Yuepeng Jiang, Qin Zhang

## Abstract

Defective decidualization is a significant pathological feature of URSA. And the potential relationship between mitochondrial fission, necroptosis and defective decidualization remains unknown. Baicalin plays an important role in regulating mitochondrial fission and programmed cell death. However, whether baicalin has a protective effect on defective decidualization in URSA has not been reported thus far. This study aims to explore the mechanisms of mitochondrial fission induced necroptosis in defective decidualization in URSA and the regulation of baicalin. First, decidual tissues were collected from URSA and health controls. And then, T-hESC was treated with lipopolysaccharide (LPS), Tyrphostin A9 (TA9), TA9+necrostatin-1(Nec-1) and TA9+baicalin during in vitro decidualization. Besides, URSA mice were established and randomly administrated with low, medium, and high doses of baicalin as well as saline. Results showed that decidualization markers prolactin (PRL) and insulin-like growth factor-binding protein-1 (IGFBP1) in patients with URSA were significantly decreased (*P*<0.05). The incidence of cell necroptosis was increased, manifested with increased Annexin V and PI positive cells, high level of pRIP3 T231(*P*<0.01) and pMLKL S358 (*P*<0.05). Moreover, mitochondrial fission was also hyperactive, featured by elevated level of Fis1 (*P*<0.01) and Drp1 (*P*<0.05). In vitro experiments, LPS was induced to trigger necroptosis of T-hESC during induced decidualization, and IGFBP1 and PRL were subsequently decreased (*P*<0.05). Besides, mitochondrial fission inducer TA9 promoted the level of necroptosis (*P*<0.05) and induced defective decidualization, which could be rescued by necroptosis inhibitor Nec-1 (*P*<0.05). In addition, baicalin could reduce mitochondrial fission (*P*<0.05), necroptosis (*P*<0.05) and ameliorate defective decidualization in vivo and in vitro (P<0.05). In conclusion, hyperactive mitochondrial fission could promote necroptosis, thus inducing defective decidualization. And baicalin could ameliorates defective decidualization in URSA by regulating mitochondrial fission induced necroptosis.

## 1. Introduction

Recurrent spontaneous abortion (RSA) is defined as two or more consecutive miscarriages before week 22 of gestation(1,2). It is one of the most distressing complications, approximately occurring in up to 5% of reproductive age couples(3). Various etiologies have been supposed to contribute to RSA, including chromosomal translocations, thrombophilias, autoimmune diseases, reproductive tract abnormality, endocrine defects, and infectious agents(4). However, 50% of cases remains elusive, namely unexplained recurrent spontaneous abortion (URSA), which poses frustrating challenges to clinicians for seeking effective strategies. Now, extensive attention and great efforts have been devoted to the study on the molecular mechanisms of URSA.

Accumulated evidence has confirmed that defective decidualization is an important pathological feature of URSA(5). Decidualization involves rapid cell proliferation, then differentiation of fibroblast-like endometrial stromal cells (ESCs) into epithelioid-like decidual stromal cells (DSCs), accompanied by extensive extracellular matrix remodeling, vascular remodeling and angiogenesis, which are crucial for embryo implantation and development. Decidualization involves profound changes in cell fate such as cell proliferation, differentiation, and apoptosis. Necroptosis, a newly discovered programmed cell death, is induced by receptor interacting serine/threonine kinase (RIP)1/RIP3 complexes, which can activate downstream mixed lineage kinase domain like pseudo kinase (MLKL), thus mediating cell death characterized by cell membrane rupture, damage-associated molecule release and inflammation(6–8), which is closely associated with various reproductive disorders such as insufficient ovarian reserve(9), pre-eclampsia(10), embryo transfer failure(11). Moreover, studies have thrown light on the correlation between necroptosis of DSCs and miscarriage and preterm birth(12). However, the mechanism connection between necroptosis and defective decidualization in URSA has not been investigated yet.

Cumulative research has focused on the central role of mitochondria in programmed cell death (PCD). Mitochondria are highly dynamic organelles constantly reshaped by fusion and fission (13). And fine balance between mitochondrial fusion and fission is crucial for cell survival and optimal functioning, while paradoxically, excessive bias of dynamic equilibrium towards fission is associated with abnormal PCD via producing oxidative stress, reducing mitochondrial membrane potential and ATP production (14). Studies have confirmed that necroptosis is mitochondria-dependent. And increased mitochondrial fission is one of the key triggers of necroptosis (13). However, whether necroptosis mediated by mitochondrial fission is related to defective decidualization in URSA has not been reported.

Baicalin (5,6,7-trihydroxyflavone, BAI) is one of the key phenolic flavonoids extracted from the root of traditional Chinese medicinal herb Scutellaria baicalensis Georgi16, 17. It has been identified that BAI exerts variou pharmacologic action, including anti-inflammatory, antioxidant and anti-cancer(15,16). Moreover, current studies have displayed that BAI is able to inhibit necroptosis(17), but whether it has a protective effect in URSA by virtue of such pharmacological activity need further in-depth studies.

In this study, we are aimed to uncover the effect of mitochondrial fission on necroptosis and decidualization in URSA and explored the effect of BAI on decidualization from its regulation on necroptosis via mitochondrial fission in URSA.

## 2. Result

### 2.1 URSA exhibited defective decidualization and increased necroptosis

F-actin distribution and the level of PRL and IGFBP1 were used to evaluate decidualization in URSA and health controls. Results verified that decidualization was defective in URSA, manifested by the spindle distribution of F-actin, and the significantly decreased mRNA level of PRL and IGFBP1 (*P*<0.05) (Figure 1A). Moreover, necroptosis in the URSA group was increased, demonstrated by increased ratio of Annexin V and PI positive cells (*P*<0.05) (Figure 1B) and decreased level of ΔΨm (*P*<0.05) (Figure 1C). Besides, pRIP3 T231(*P*<0.05) and pMLKL S358 (*P*<0.01) in the decidua tissue of URSA were significantly increased (Figure 1D, 1E). These results suggested that URSA exhibited defective decidualization, accompanied by increased cell necroptosis in decidua tissue.

**Figure 1.**
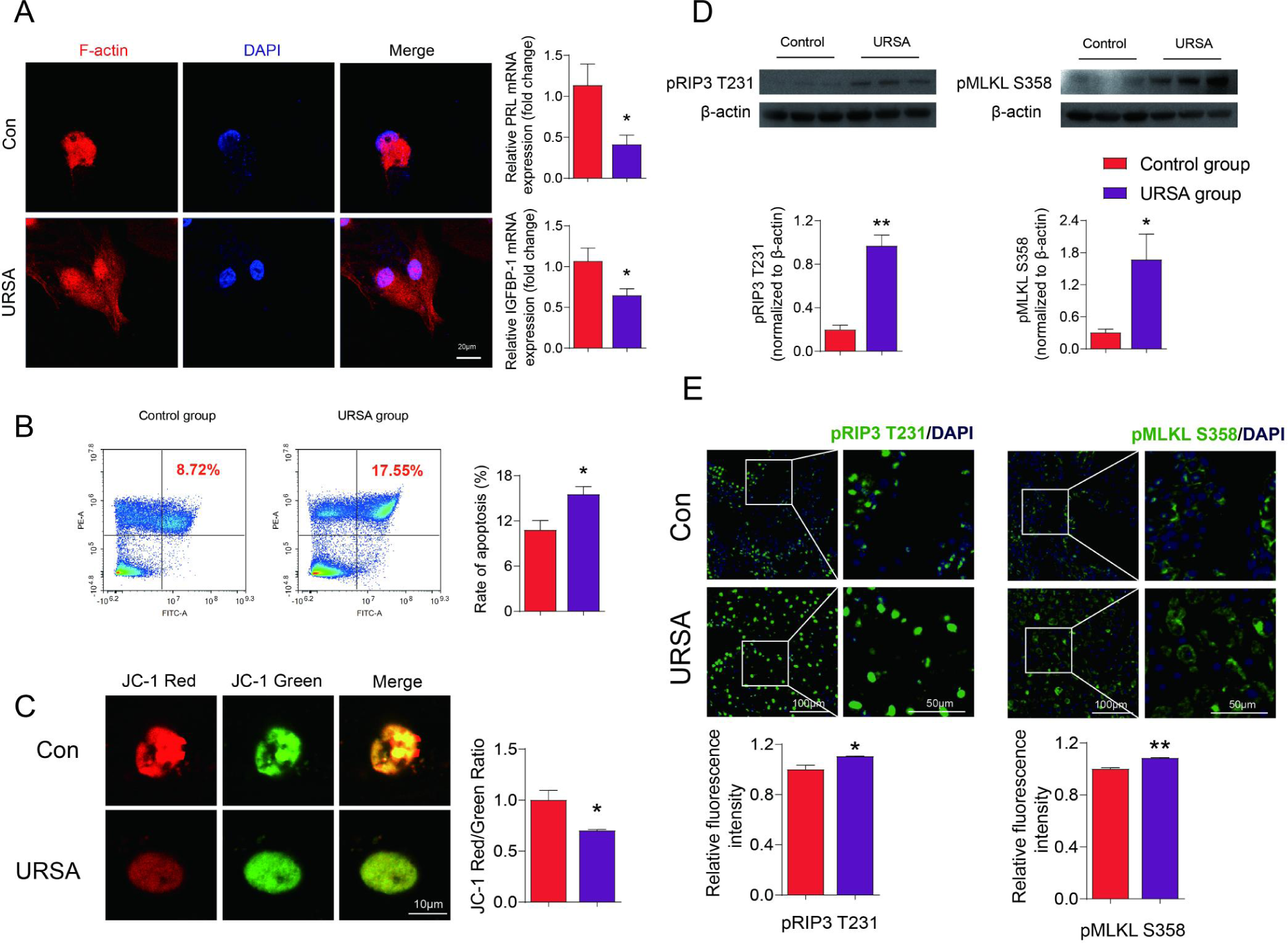
Decidual tissue derived from URSA featured with defective decidualization and increased level of necroptosis A. Human F-actin, PRL and IGFBP1 in different groups. B. The number of Annexin V and PI positive cells. C. The ratio of Red/Green fluorescence intensity. D. The levels of pRIP3 T231 and pMLKL S358 of Western blot. E.Immunofluorescence levels of pRIP3 T231 and pMLKL S358. * P<0.05 vs. control group, ** P<0.01 vs. control group. By repeated-measures one-way ANOVA followed by post hoc Dunnett’s multiple comparisons test. In this and subsequent figures, the experimenters were blinded to the treatment condition(s).

### 2.2 LPS induced necroptosis could led to defective decidualization in vitro

As necroptosis were significantly increased in URSA, we further explored the effect of necroptosis on decidualization. Thus, LPS was utilized to induced necroptosis during in vitro decidualization (18,19). The results showed that the ratio of JC-1 red/green fluorescence intensity in LPS group was reduced (*P*<0.05) (Figure 2A), accompanied by an increase in the levels of pRIP3 T231 and pMLKL S358 in LPS group (*P*<0.05) (Figure 2B-C). Besides, the mRNA levels of PRL and IGFBP1 were synchronously decreased in LPS group (*P*<0.05) (Figure 2D-E). These results indicated that necroptosis was a key cellular event leading to defective decidualization in URSA.

**Figure 2.**
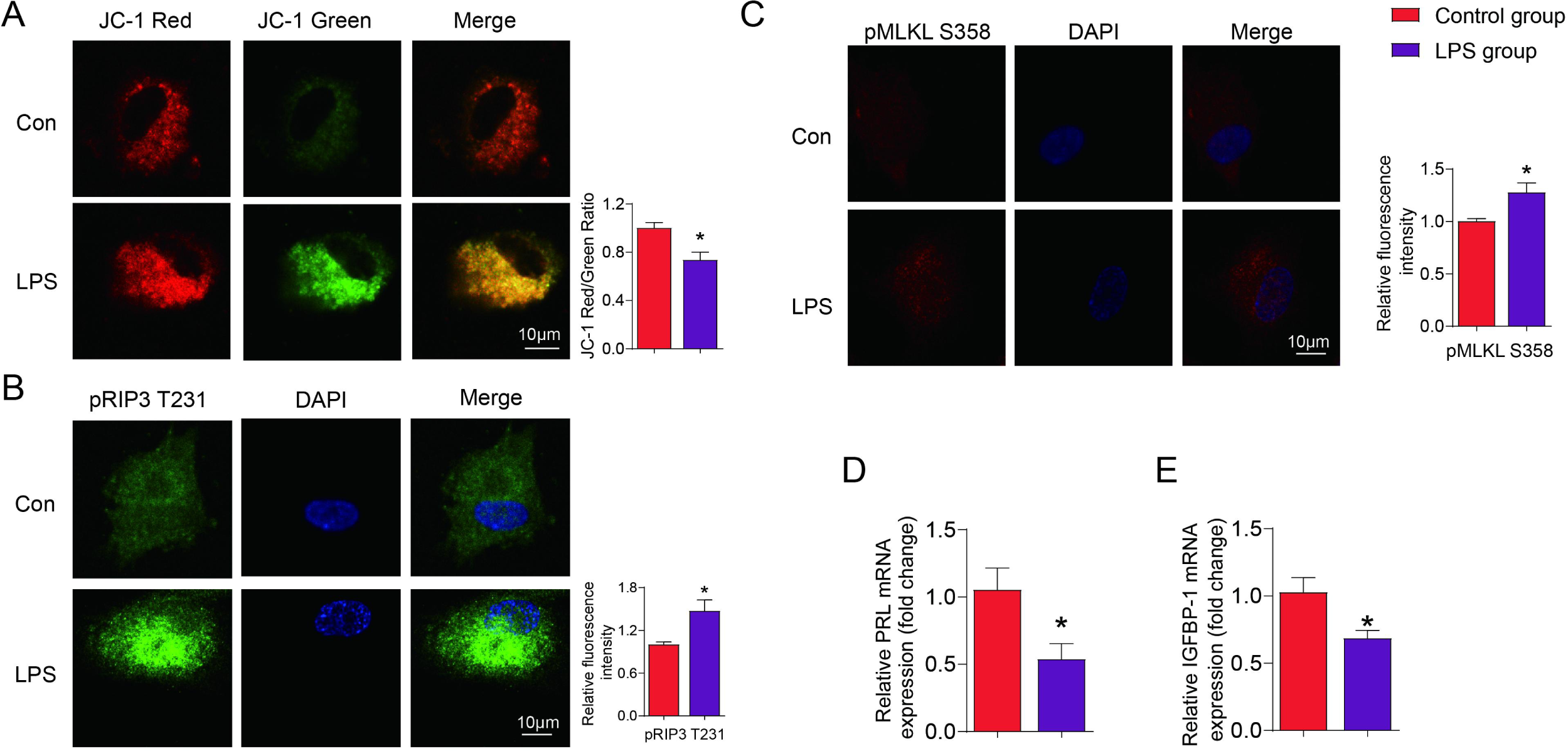
LPS induced necroptosis in vitro decidualization A. The ratio of red/green fluorescence intensity. Immunofluorescence intense of pRIP3 T231 (B) and pMLKL S358 (C). The mRNA levels of PRL (D) and IGFBP1(E). * P<0.05 vs. control group, ** P<0.01 vs. control group. By repeated-measures one-way ANOVA followed by post hoc Dunnett’s multiple comparisons test.

### 2.3 Mitochondrial fission was significantly active in URSA

Considering that mitochondrial fission is closely related to programmed cell death, we further explored the correlation between mitochondrial fission and cell necroptosis in URSA. Studies have shown that mitochondrial fission protein 1 (Fis1) can promote mitochondrial fission by promoting the recruitment of Drp1 to mitochondria(20). Given their pivotal role in mitochondrial fission, we further detected the level of Drp1 and Fis1 in URSA and health controls. The results displayed that the expression profiles of Drp1 (*P*<0.01) and Fis1 (*P*<0.05) detected by Western blot were substantially increased (Figure 3A-B), which was consistent with the immunofluorescence assay (Drp1 and Fis1: *P*<0.05) (Figure 3C-D). This indicated that mitochondrial fission was significantly active in URSA.

**Figure 3.**
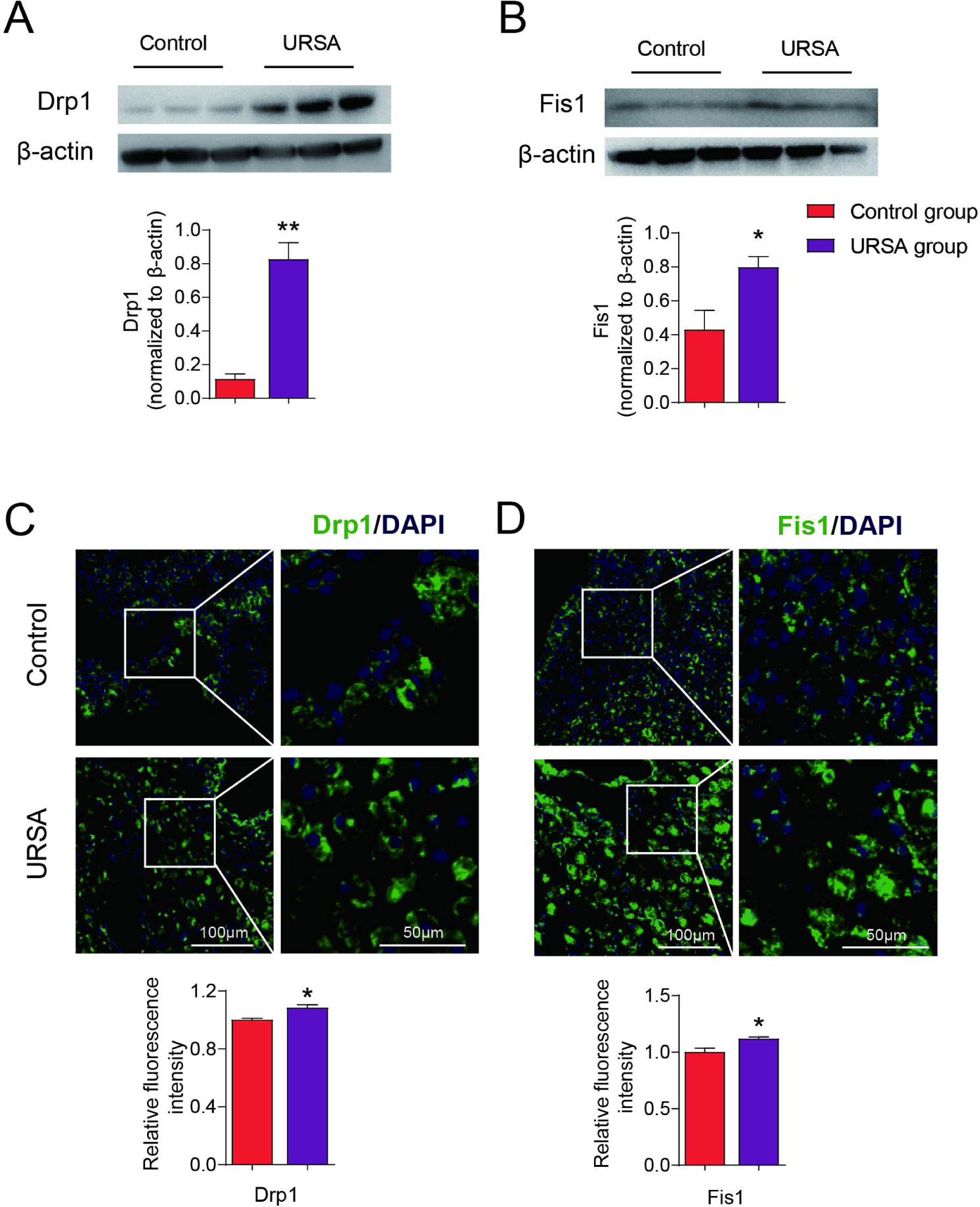
Mitochondrial fission was induced in URSA A. Human Drp1 levels with Western blot in different groups. B. Human Fist1 levels with Western blot in different groups. C. Immunofluorescence levels of Drp1. D. Immunofluorescence levels of Fist1. * P<0.05 vs. control group, ** P<0.01 vs. control group. By repeated-measures one-way ANOVA followed by post hoc Dunnett’s multiple comparisons test.

### 2.4 Mitochondrial fission could induce necroptosis and defective decidualization in vitro

Moreover, given that mitochondrial fission is increased in URSA, and that mitochondrial fission is closely related to necroptosis(21–23), we further explored the effect of mitochondrial fission on necroptosis and decidualization in vitro. First, TA9 was applied to successfully induce mitochondrial fission, characterized by the increased expression of (western blot: Drp1 *P*<0.05; Fis1 *P*<0.01. immunofluorescence: Drp1 and Fis1 *P*<0.05) (Figure 4A-D). And western blot showed that the level of pRIP3 T231 and pMLKL S358 were also increased in TA9 group (*P*<0.05) (Figure 5A-B), which was consistent with the immunofluorescence assay (P<0.01) (Figure 5C-D). Besides, TA9 significantly resulted in decidualization defects, with observably decreased levels of PRL (*P*<0.05) and IGFBP1 mRNA (*P*<0.01) (Figure 5E-F). Moreover, The effect of TA9 on decidualization could be mitigated by necroptosis inhibitors, Nec-1, which act as a specific allosteric inhibitor of receptor interacting protein-1(RIP1) kinase, a downstream signaling molecule of the death receptor induced non-necroptosis cell death pathways(24–26). As shown in Figure 5, the levels of pRIP3 T231 (*P*<0.05) and pMLKL S358 (*P*<0.05) were reduced in the TA9+Nec-1 group when compare with the TA9 group (Figure 5A-D), and the mRNA levels of IGFBP1 and PRL were also rescued (P<0.05) (Figure 5E-F). The above experiments suggested that over activated mitochondrial fission could mediate defective decidualization via promoting necroptosis.

**Figure 4.**
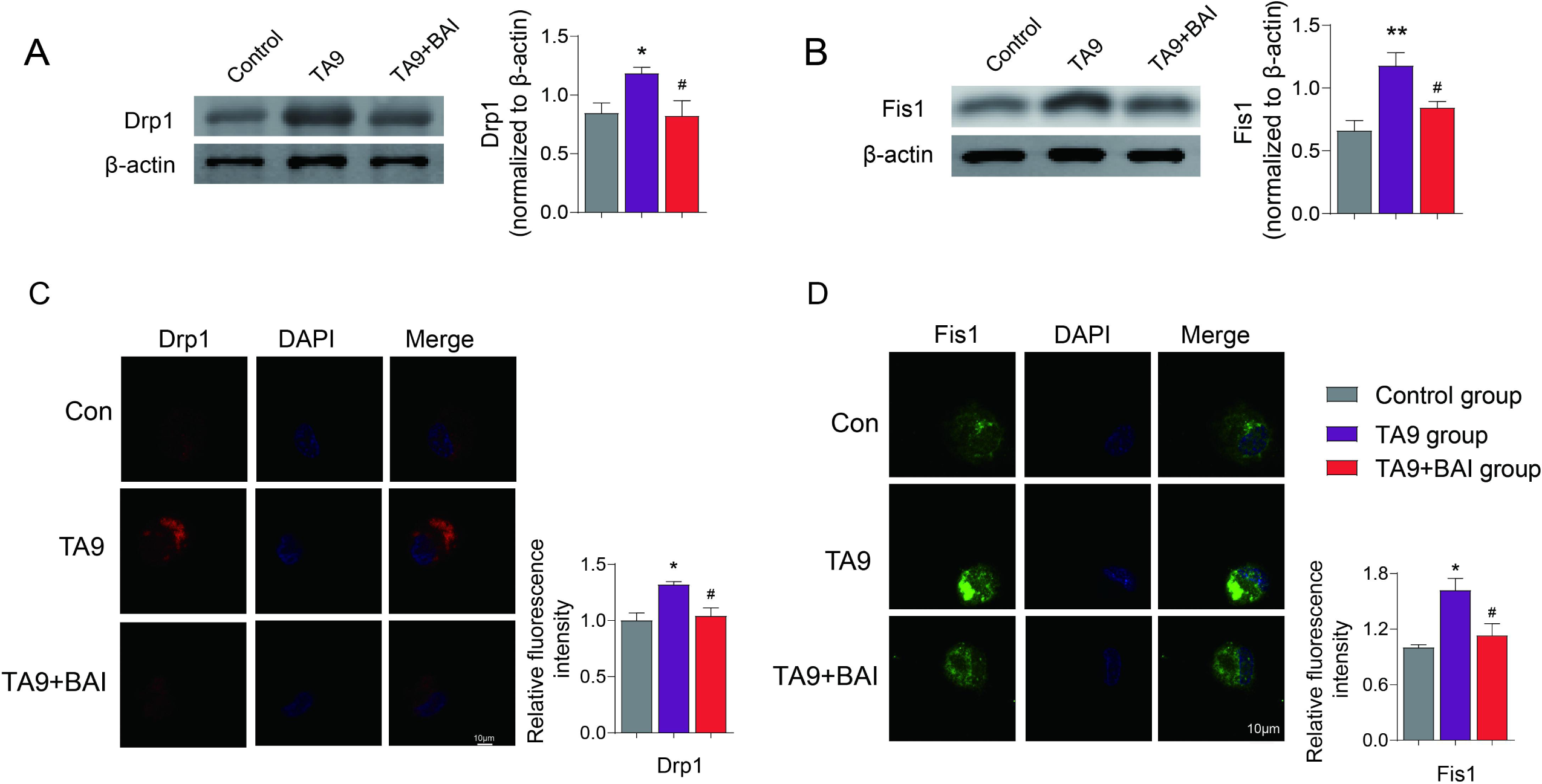
TA9 induced mitochondrial fission A. The levels of Drp1 with Western blot in three groups in vitro. B. The levels of Fis1 with Western blot in three groups. C. Immunofluorescence levels of Drp1. D. Immunofluorescence levels of Fis1. * P<0.05 vs. control group, ** P<0.01 vs. control group. ^#^ P<0.05 vs. TA9 group. By repeated-measures one-way ANOVA followed by post hoc Dunnett’s multiple comparisons test.

**Figure 5.**
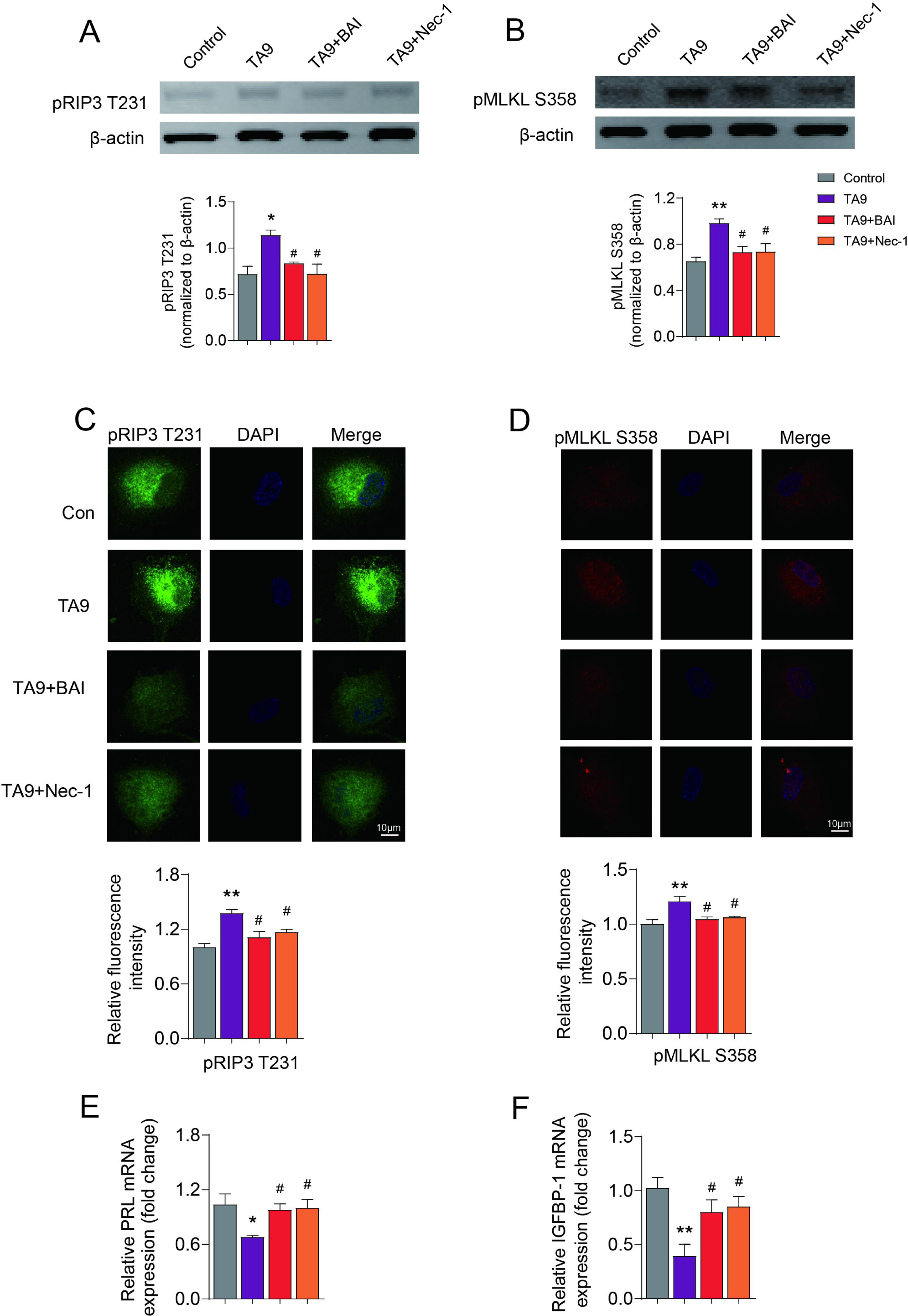
Baicalin could improve decidualization by affecting the necroptosis A. The levels of pRIP3 T231 with western blot. B. The levels of pMLKL S358 with western blot. The fluorescence intensity of pRIP3 T231 (C) and pMLKL S358 (D). E. The mRNA levels of PRL. F. The mRNA levels of IGFBP-1. * P<0.05 vs. control group, ** P<0.01 vs. control group. ^#^ P<0.05 vs. TA9 group. By repeated-measures one-way ANOVA followed by post hoc Dunnett’s multiple comparisons test.

### 2.5 Baicalin could improve defective decidualization by affecting necroptosis triggered by mitochondrial fission in URSA

In this part, we investigated the effect of BAI on decidualization from the aspect of mitochondrial fission induced necroptosis. And the results confirmed that baicalin treatment could reduce the intensity of ROS in mitochondrial and elevate the ratio of JC-1 Red/Green fluorescence intensity (*P*<0.05) (Figure 6A-B). Besides, BAI could inhibit the expression of Drp1 and Fis1 triggered by TA9 (*P*<0.05) (Figure 4A-D), reduced the level of pRIP3 T231 (*P*<0.05) and pMLKL S358 (*P*<0.05) (Figure 5A-D), and improve the mRNA level of PRL (P<0.05) and IGFBP1 (*P*<0.05) (Figure 5E-F).

**Figure 6.**
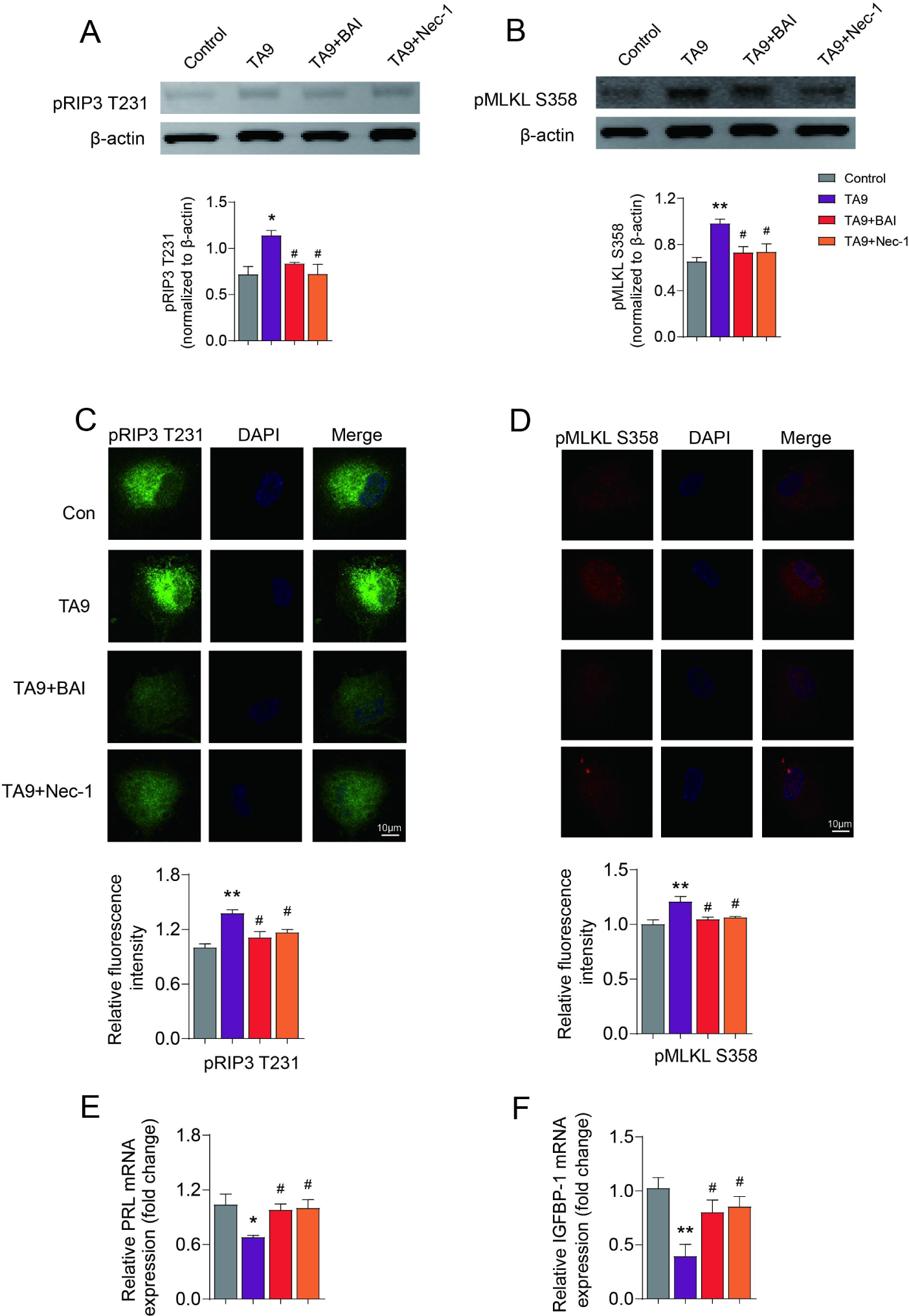
Baicalin could impact mitochondrial fission A. The fluorescence intensity of ROS in mitochondrial. B. The intensity of JC-1 red/green ratio. * P<0.05 vs. control group, ** P<0.01 vs. control group. ^#^ P<0.05 vs. TA9 group. By repeated-measures one-way ANOVA followed by post hoc Dunnett’s multiple comparisons test.

In addition, we further investigated that the intervention of baicalin on URSA mice. The results showed that baicalin reduced the expression of Drp1 (*P*<0.05), Fis1 (*P*<0.01), pRIP3 T231 (*P*<0.05) and pMLKL S358 (*P*<0.05) (Figure 7A-D), increased the mRNA levels of PRL and IGFBP1 (*P*<0.05) in a dose-dependent manner (Figure 7E-F). Besides, baicalin significantly reduced embryo absorption rate in a dose-dependent manner on day 8.5 (*P*<0.05) and day 14.5 of gestation(*P*<0.01), respectively (Figure 7G-H). Combined with the vitro and vivo experiments, we demonstrated that baicalin could improve decidualization by affecting the necroptosis triggered by mitochondrial fission in URSA.

**Figure 7.**
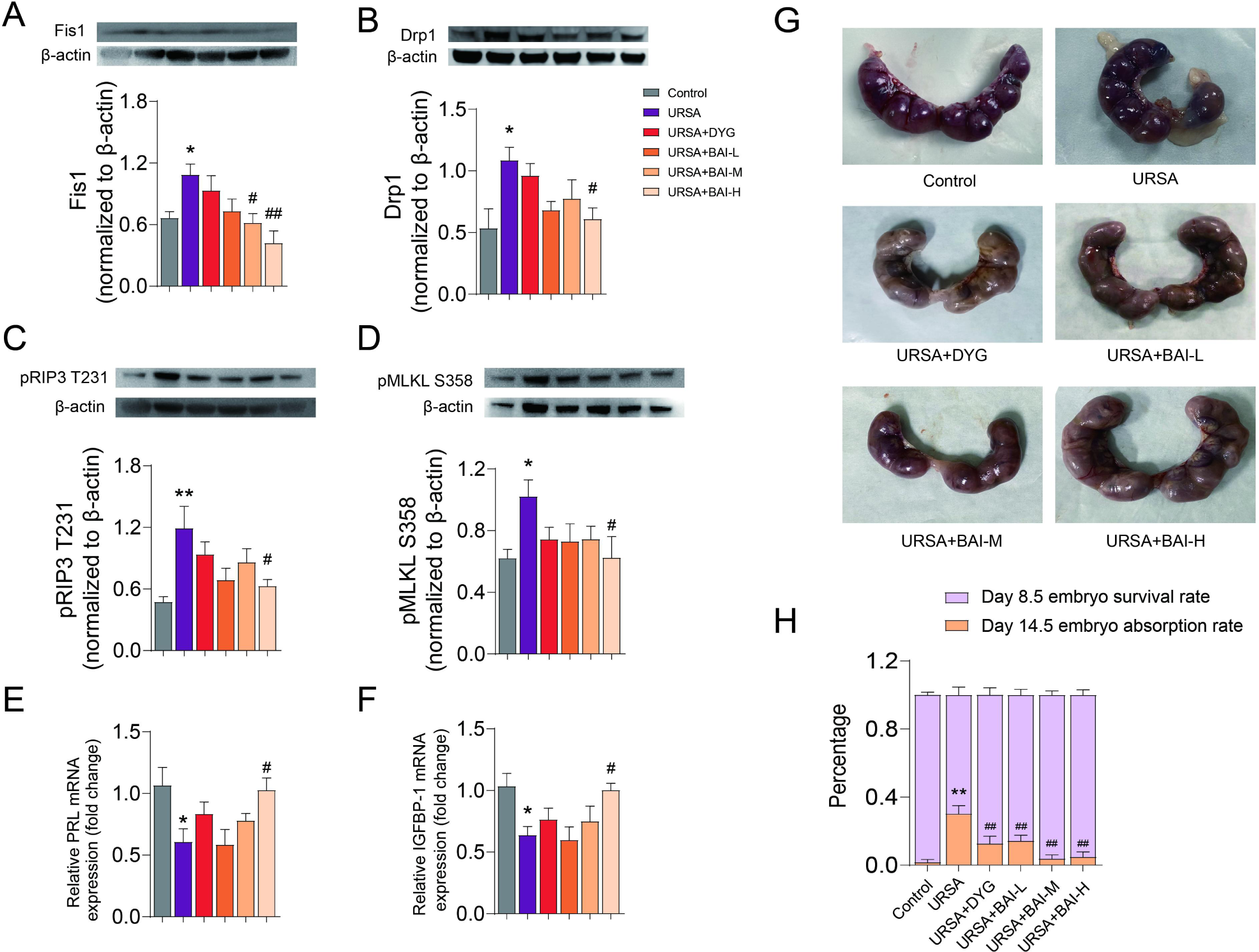
Baicalin could improve decidualization by affecting the necroptosis triggered by mitochondrial fission in URSA mice A. Drp1 Western blot expression in URSA mice groups. B. Fis1 Western blot expression. The pRIP3 T231 (C) and pMLKL S358 (D) levels with Western blot. Expression levels of PRL (E) and IGFBP-1 (F) detected by qRT-PCR. G. The mice uterus with embryos on 14.5 day of pregnancy. H. The embryo survival rate on 8.5 day and the embryo absorption rate on 14.5 day. * P<0.05 vs. control group, ** P<0.01 vs. control group. ^#^ P<0.05 vs. URSA group. By repeated-measures one-way ANOVA followed by post hoc Dunnett’s multiple comparisons test.

## 3. Discussion

In this study, we uncovered increased mitochondrial fission, enhanced necroptosis, and defective decidualization in the decidual tissue of URSA, and then we demonstrated in vitro that hyperactivated mitochondrial fission promoted necroptosis and defective decidualization, and the defective decidualization caused by mitochondrial fission can be rescued when necroptosis was inhibited. Thus, it could be deduced that abnormally active mitochondrial fission was able to trigger necroptosis, which in turn mediated the defective decidualization of URSA. In addition, we demonstrated in vivo and in vitro that baicalin was capable of improving decidualization in URSA by regulating mitochondrial fission induced necroptosis, thereby preventing recurrent embryo loss.

In mice, embryo implantation occurs around midnight on day 4 of pregnancy(27). For this process to proceed successfully, decidualization occur induced by steroid hormone, accompanied by proliferation, differentiation and PCD, which renders it competent for embryo attachment(28). Necroptosis is a relatively new concept of non-apoptotic programmed cell death that can be triggered by exogenous bacterial toxins or endogenous cellular senescence(29). Similar to apoptosis, necrosis is initiated by tumor necrosis factor alpha (TNF-α), leading to the activation of receptor-interacting protein kinase 1 (RIP1)(30). RIP3 is then recruited to RIP1 to form a necrosome that catalyzes phosphorylation of the mixed-lineage kinase domain-like (MLKL). After that, pMLKL S358 drills the cellular membrane and directly disrupts membrane integrity and triggers necrosis(31). Necroptosis have been found that related to overactive inflammation, which is a threat to maternal vascular adaptation and trophoblast invasion, thus affecting embryo implantation and development(32). Besides, hyperactivated necroptosis has been verified to be a contributing factor in abnormal pregnancy(33,34). However, the association between necroptosis and URSA has not been revealed so far. Our study revealed a significantly increased level of necroptosis in decidual tissue both in women with URSA and in URSA mice model. We speculated that excessive necroptosis destroyed the biological function of DSCs and stimulates an overly aggressive inflammatory response, thus resulting in defects of decidualization, which fail to provide a conducive maternal environment for implantation and embryo development. It is worth mentioning that this is the first time that we have revealed the effect of necroptosis on URSA.

More importantly, we also identified abnormal mitochondrial dynamics as a possible mechanism for triggering necroptosis. Mitochondria are highly dynamic organelles reshaping by constant fission and fusion, which are regarded as central regulator of the decision between cellular survival and demise. Mitochondrial fission is regulated by DRP1 and its receptors, including mitochondrial fission factor (MFF), FIS1, mitochondrial dynamics protein of 49 kDa (MID49) and mitochondrial dynamics protein of 51 kDa (MID51). Following fission, the smaller clusters may become nearly spherical with diameters of about a few hundred nanometers, which helps to remove dysfunctional mitochondria from cardiomyocytes. However, if fission is not controlled and balanced by fusion, the network becomes too fragmented which can result in glucose oxidation, mitochondrial inner membrane potential decline, and hence the downregulation of ATP and necroptosis(25). Studies have confirmed that mitochondrial fission was involved in necroptosis(24,26). For instance, Lipopolysaccharide was proved to trigger drp1-mediated mitochondrial fragmentation leading to necrosis in murine primary macrophages. Drp1 inhibition reduces hydrogen peroxide (H2O2)-induced necrosis(35). Interestingly, various studies have reported that the necroptosis-related proteins are located at mitochondria and interact with Drp125, Wang et al. found that mitochondrial damage preceded necroptosis occurrence and RIPK1/RIPK3 complex directly phosphorylated Drp1 and triggered its translocation to mitochondria2(36,37). These studies suggest that mitochondrial fission, regulated by Drp1, might be an intermediate event of necroptosis. At present, the influence of mitochondrial dynamics on the endometrial cycle has been confirmed. Research showed that the decidualized hESC have shorter mitochondria than those in the proliferative phase, which suggested that the deciduating phase might be accompanied by an increase in mitochondrial fission(38). However, the effect of unbalanced mitochondrial fission on decidualization has not been reported. In our study, we first verified the excessive mitochondrial fission in decidual tissue in URSA and triggering mitochondrial fission could induce defective decidualization by provoking necroptosis. Studies have shown that excessive mitochondrial fission is associated with reactive oxygen species (ROS) leakage(39), which is the major signal of triggering necroptosis. After fission, mitochondria cannot to acidic lysosomes for circulating, and continuous overproduction of ROS in mitochondrial (mtROS) eventually leads to necroptosis(22). Thus, we speculate that excessive mitochondrial fission in URSA may increase necroptosis activated by mtROS, thereby disrupting decidualization.

Baicalin as a bioactive component present in Chinese herbal medicine, possesses a wide range of pharmacological activities with anti-apoptosis capability**(40)**. Besides, A potent protective effect of baicalin has been discovered on various disease by regulating mitochondria-related apoptosis**(41)**. Furthermore, baicalin is able to exert a anti-necroptosis**(42)** effects via attenuating the expression profiles of Drp1**(43)**. Besides,it has been identified that baicalin is capable of attenuating acrolein induced elevation in RIPK-3 which executes the necroptosis**(44)**. However, the interference of baicalin on mitochondrial fission induced necroptosis in URSA has not been reported. Here we suggested a novel point that baicalin conspicuously improve decidualization defects and reduced miscarriage by inhibiting the expression of Drp1 and Fis1, and then suppressed mitochondrial fission, and reduced necroptosis, this properties of baicalin could significantly attenuate defective decidualization caused by necroptosis.

## 4. Conclusion

Our study showed that mitochondrial fission was increased in the URSA group, and activation of mitochondrial fission promoted necroptosis and thus disrupted decidualization and lead to recurrent miscarriage. Besides, baicalin was capable of improving decidualization in URSA by regulating mitochondrial fission and necroptosis, thereby preventing embryo loss (Figure 8). In conclusion, our study is the first to clarify the role of necroptosis in defective decidualization of URSA, and to clarify the molecular mechanism of mitochondrial dynamics triggering necroptosis to participate in defective decidualization. In addition, we also clarify the potential mechanism of baicalin treatment of URSA. The above results provide a new perspective on the understanding of the pathological mechanism of URSA and provide information for targeted treatment strategies for URSA.

**Figure 8.**
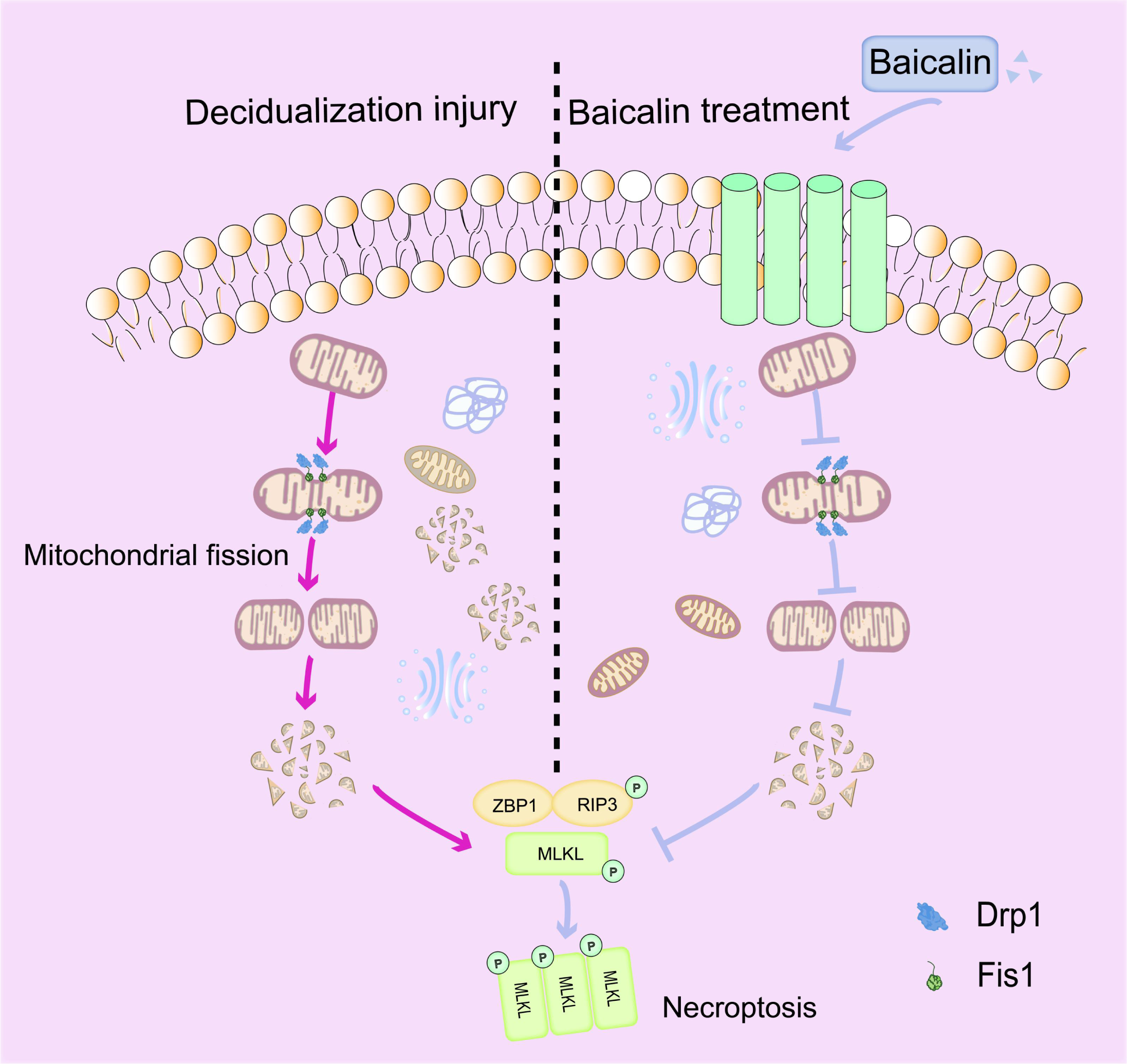
Mechanism hypothesis diagram

## 5. MATERIALS AND METHODS

### 5.1 Reagents and antibodies

Fetal bovine serum (FBS) (Cat. 10099141; Gibco), Penicillin (Cat. ST488-1; Beyotime), Streptomycin (Cat. ST488-2; Beyotime), 8-Br-cAMP (Cat. B5386; Sigma), MPA (Cat. B1510; APExBIO), mitochondrial fission inhibitor: Mdivi-1 (Cat. SC8028; Beyotime), Tyrphostin A9 (Cat. HY-15511; MCE), baicalin (Cat. 572667; Sigma-Aldrich). mitochondrial membrane potential assay kit with JC-1 (Cat. C2006; Beyotime), mitoSOX red mitochondrial superoxide indicator (Cat. LX3608; Warbio), coraLite®594-Phalloidin (red) (Cat. PF00003; proteintech). Western blot experiments were performed using the following antibodies: rabbit anti-Drp1 monoclonal antibody (1:1000; Cat. 8570; CST), mouse anti-Fis1 monoclonal antibody (1:1000; Cat. sc-376447; SANTA), rabbit anti-Phospho-MLKL-T357/S358/S360 polyclonal antibody (1:1000; Cat. AP0949; ABclonal), rabbit anti-Phospho-RIP3-T231/S232 polyclonal antibody (1:1000; Cat. AP1260; ABclonal), rabbit anti-β-Actin monoclonal antibody (1:1000; Cat. 4970; CST), goat anti-rabbit IgG (H&L) (1:5000; Cat. 926-32211; LI-COR). Immunofluorescence experiments were performed using the following antibodies: rabbit anti-TOM20 antibody (1:200; Cat. 42406; CST), mouse anti-ICAM1 antibody (1:200; Cat. Ab171123; Abcam), goat anti-rabbit IgG H&L (Alexa Fluor® 488) (1:200; Cat. ab150077; Abcam), goat anti-rabbit IgG H&L (Cy3®) (1:200; Cat. ab6939; Abcam), goat anti-rabbit IgG H&L (Alexa Fluor® 647) (1:200; Cat. Ab150083; Abcam), goat anti-mouse IgG H&L (Alexa Fluor® 488) (1:200; Cat. ab150113; Abcam), goat anti-mouse IgG H&L (Alexa Fluor® 647) (1:200; Cat. ab150115; Abcam).

### 5.2 Tissue samples

The study aimed to investigate the potential differences in decidua tissues between healthy women and patients with URSA. Decidua tissues were collected from both groups of participants who underwent vacuum aspiration to terminate their pregnancies during the first trimester of gestation at the Hangzhou Hospital of Traditional Chinese Medicine Affiliated to Zhejiang Chinese Medicine University. The URSA group included women who had experienced at least two miscarriages, excluding definite laboratory detectable etiologies(45), while the control group consisted of healthy women who had given birth to at least one healthy child and had no history of adverse pregnancy (terminated for nonmedical reasons) (45). After the surgical procedure, decidual tissue specimens were collected and prepared for further experiments. The study protocol was approved by the Ethics Committee of the Hangzhou Hospital of Traditional Chinese Medicine Affiliated to Zhejiang Chinese Medicine University, and the research was conducted in accordance with the principles of the Declaration of Helsinki. Informed consents were collected from all the subjects, indicating that they were aware of the purpose of the study, the potential risks and benefits, and their rights as participants. The collection and use of human decidua tissues for research purposes were carried out with strict adherence to ethical guidelines and regulations to ensure the protection of human subjects.

### 5.3 URSA mouse model establishment and drug treatment

In this study, adult female CBA/J, male DBA/2, and male BALB/c mice were used. These mice were purchased from Beijing Huafukang Biotechnology Co. Ltd. (China) and were kept under standard laboratory conditions, including a temperature of 19-23 °C, 12-hour light/dark cycles, and 40-60% humidity. After a period of adaptation, female CBA/J mice were mated with either male BALB/c or male DBA/2 mice at a 2:1 ratio to construct mouse models of normal pregnancy or of URSA. The URSA mice were randomly divided into five groups: URSA group, Dydrogesterone group, and low, medium, and high dose groups of baicalin (10 mg/kg, 20 mg/kg or 50 mg/kg per day). The oral administration in each group started from 0.5 days of gestation, marked by the presence of a postcoital vaginal plug. The decidual tissues of the mice were collected at 8.5 or 14.5 days of gestation. All animal experiments were conducted in compliance with the LARC Animal Ethics Code and the ethics number for the study was 20190909-07.

### 5.4 Cell culture and in vitro decidualization of T-hESC

In accordance with prior literature(46), human telomerase reverse transcriptase-immortalized human endometrial stromal cells (T-hESCs, ATCC, CRL-4003) were cultured in phenol red-free DMEM/F12 medium supplemented with 10% charcoal-stripped fetal bovine serum (CS-FBS), 100U/mL penicillin, and 100μg/mL streptomycin. Decidualization was induced in vitro by adding 0.5mM 8-Br-cAMP and 1μM MPA to the culture medium, and the induction process was carried out over a period of 6 days, with the medium being changed every 2 days. The decidualization of T-hESCs was assessed by evaluating the morphological phenotype and measuring the levels of decidual markers, including PRL mRNA and IGFBP1 mRNA (47).

### 5.5 Cell processing and grouping

T-hESCs were divided randomly into several treatment groups: DSC group, TA9 group, baicalin group, necrostatin-1 (Nec-1) group, and lipopolysaccharide (LPS) group, in which TA9 served as an inducer of mitochondrial fission, Nec-1 as a necroptosis inhibitor, and LPS as an endotoxin to induce necroptosis(48). The DSC group was treated with 1μM MPA and 0.5mM 8-Br-camp for 6 days to induce decidualization. The TA9 group was treated with 1μM TA9 for 48 hours on the fourth day of decidualization to induce mitochondrial fission. The baicalin group was treated with 1μM TA9 and baicalin (50μM) on the fourth day of decidualization for 24h(49). The Nec-1 group was treated with 1μM TA9 and 40µM Nec-1 for 48 hours on the fourth day of decidualization to inhibit necroptosis. The LPS group was treated with 10μg/mL LPS for 24 hours during decidualization to induce necroptosis.

### 5.6 Quantitative real-time reverse transcription polymerase chain reaction (qRT-PCR)

Quantitative real-time PCR analysis was conducted following previously described protocols. Total RNA from cells was extracted using TRIzol reagent (Cat. 15596026; Invitrogen), and reverse transcription was performed using the HiFiScript cDNA Synthesis Kit (Cat. CW2569M; CWBIO). The resulting cDNA was subjected to quantitative real-time PCR using the SYBR Green kit (Solarbio) on an Exicycler TM 96 instrument (Bioneer) with β-actin serving as the reference gene. The relative mRNA expression level was calculated using the comparative 2−ΔΔCt method based on the threshold cycle (Ct). The primer sequences used for real-time PCR are provided in Table 1.

**Table 1.**
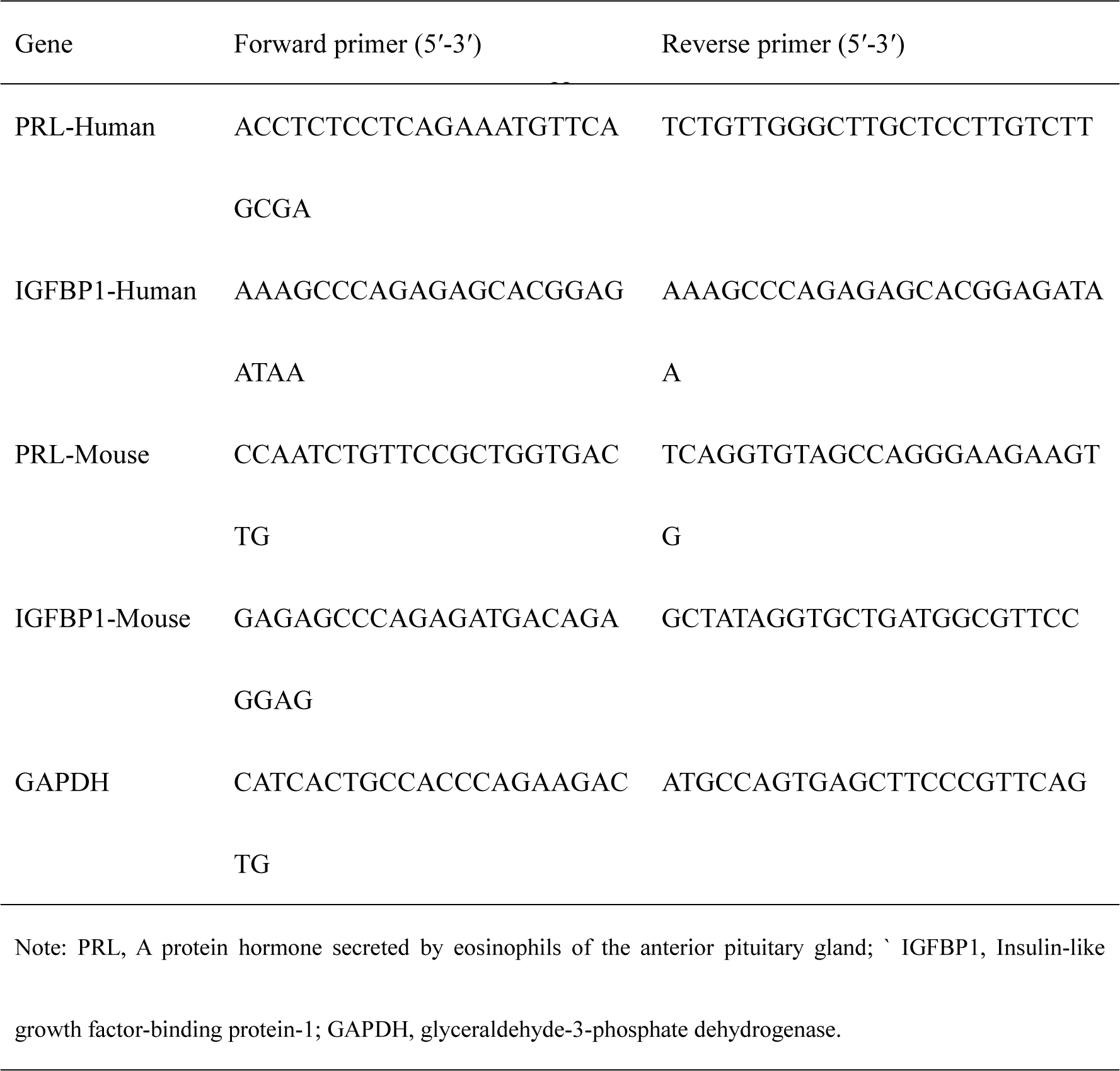
Primer used for qRT-PCR

### 5.7 Flow cytometry

The endometrial tissue from women was chopped, washed with Hank’s balanced salt solution (HBSS), and digested with 0.15% Collagenase I (Sigma-Aldrich, USA) at 37 °C for 30 minutes. Subsequently, the ESCs were separated from the glandular tissue by filtering the digested tissue through a 30μm screen. The cells were then resuspended with PBS and enumerated. A total of 1×106 cells were centrifuged at 1000 r/min for 5 minutes, and the supernatant was discarded. The cells were gently resuspended with 500μL Annexin V-FITC binding solution, and 5μL Annexin V-FITC was added, mixed gently, and incubated in the dark at room temperature for 15 minutes. The cells were then centrifuged at 1000 r/min for 5 minutes, the supernatant was discarded, and 500μL Annexin V-FITC binding solution was added to gently resuspend the cells. Finally, 10μL propidium iodide (PI) staining solution was added, mixed gently, and the cells were placed in an ice bath for 10 minutes in the dark. The flow cytometry (Beckman Coulter, USA) was used to detect the apoptosis rates, and FlowJo 7.6.2 software (BD, USA) was employed for data analysis.

### 5.8 Immunofluorescence

The T-hESCs were fixed in 4% paraformaldehyde solution in PBS for 200 minutes, followed by permeabilization with 0.3% Triton X-100 for 15 minutes. After blocking with 5% bovine serum albumin (BSA) for 30 minutes, the cells were incubated overnight with primary antibodies: rabbit anti-TOM20 and mouse anti-ICAM1 (both diluted 1:200). Subsequently, the cells were incubated with secondary antibodies, including goat anti-rabbit IgG H&L (Cy3) and goat anti-mouse IgG H&L (Alexa Fluor® 488) (both diluted 1:200), for 2 hours. Phalloidin was then added and incubated for 20 minutes. Images were acquired using a laser confocal microscope (LSM880, Zeiss).

### 5.9 Western blotting

Initially, RIPA lysis buffer (Cat. P0013B; Beyotime Biotechnology), supplemented with protease inhibitors (Cat. B14002; Bimake), phosphatase inhibitors (Cat. B15002; Bimake), and pierce^TM^ universal nuclease for cell lysis (Cat. 88700; Thermo Scientific) was utilized to extract total protein from human and mouse decidual tissue or T-hESCs. The protein concentration was determined using a BCA assay, and 40 µg/lane of protein was loaded onto 12.5% SDS-PAGE gels and transferred onto PVDF membranes. After blocking with 10% skim milk at room temperature for 1 hour, the membranes were incubated overnight at 4 °C with the following primary antibodies: rabbit anti-Drp1 monoclonal antibody (1:1000; Cat. 8570; CST), mouse anti-Fis1 monoclonal antibody (1:1000; Cat. sc-376447; SANTA), rabbit anti-Phospho-MLKL-T357/S358/S360 polyclonal antibody (1:1000; Cat. AP0949; ABclonal), rabbit anti-Phospho-RIP3-T231/S232 polyclonal antibody (1:1000; Cat. AP1260; ABclonal), and rabbit anti-β-Actin monoclonal antibody (1:1000; Cat. 4970; CST). Subsequently, the membranes were incubated with goat anti-rabbit IgG (H&L) (1:5000; Cat. 926-32211; LI-COR) and goat anti-mouse IgG (H&L) (1:5000; Cat. 926-68070; LI-COR) secondary antibodies for 2 hours. Quantitative analysis was performed by Image J software after scanning the membranes using an Odyssey fluorescence imaging system.

### 5.10 Mitochondrial membrane potential analysis

Mitochondrial membrane potential ΔΨm was assessed using a JC-1 fluorescent probe. Upon high mitochondrial membrane potential, JC-1 formed aggregates in the matrix of mitochondria and exhibited red fluorescence. Conversely, under low mitochondrial membrane potential, JC-1 was present as a monomer in the matrix of mitochondria and generated green fluorescence. Following the addition of 500μL of JC-1 staining working solution under light-proof conditions at 37°C with 5% CO_2_ for 20 min, cells from different groups were scraped off with a spatula and resuspended in 300μL PBS. Image acquisition was performed using a laser confocal microscope, and fluorescence intensity was quantified using Image J.

### 5.11 Measurement of mitochondrial ROS production

We utilized MitoSOX Red Mitochondrial Superoxide Indicator (Cat. LX3608; Warbio) to quantify the levels of reactive oxygen species (ROS). Cells were seeded onto 24-well cell culture plates and incubated for 24 hours before being treated with 5μM of MitoSOX Red Mitochondrial Superoxide Indicator at 37°C for 10 min in the dark, as per the manufacturer’s instructions. The cells were subsequently washed thrice with PBS and counterstained with DAPI for 10 min. Finally, we measured the fluorescence intensity at 647nm (red fluorescence) using a confocal laser-scanning microscope.

### 5.12 Statistical analyses

The statistical analysis was carried out using GraphPad Prism 8 (San Diego, California, USA). Differences between groups were evaluated using either Student’s t-test or one-way analysis of variance (ANOVA), followed by Dunnett’s post-hoc test. All data are presented as mean ± standard deviation (Mean ± SD) and were obtained from at least three independent experimental replicates. A value of P<0.05 was considered to be statistically significant.

## Acknowledgments

Not applicable.

## Authors contribution

X.X.Z and Y.Z performed the experiments; Q.J.Y, Y.P.J, S.X.W and Q.Z conceived and designed the experiments; J.M, Y.Z, and Y.Z analyzed the data, X.X.Z and Y.Z wrote the paper. All authors contributed to manuscript preparation.

## Institutional Review Board Statement

The study was conducted according to the guidelines of the Declaration of Helsinki and approved by the Laboratory Animal Center of Zhejiang Chinese Medical University (Ethics number: IACUC-20220913-12).

## Informed Consent Statement

Not applicable.

## Data availability statement

The original contributions presented in the study are included in the article/supplementary material, further inquiries can be directed to the corresponding authors.

## Abbreviations

BSA: bovine serum albumin
Drp1: Dynamin-related protein 1
DSCs: decidual stromal cells
ESCs: endometrial stromal cells
FBS: Fetal bovine serum
Fis1: Mitochondrial fission 1 protein
HBSS: Hank’s balanced salt solution
IGFBP1: Insulin-like growth factor-binding protein-1
LPS: Lipopolysaccharide
MFF: mitochondrial fission factor
Nec-1: Necrostatin-1
PI: propidium iodide
pMLKLS358: Mixed lineage kinase domain-like protein S358
pRIP3T231: Receptor-interacting protein 3 T231
PRL: Prolactin
ROS: reactive oxygen species
mtROS: ROS in mitochondrial
TA9: Tyrphostin A9
TNF-α: tumor necrosis factor alpha
T-hESC: Transcriptase-immortalized human endometrial stromal cells
URSA: Unexplained Recurrent Spontaneous Abortion
H2O2: hydrogen peroxide

## Reference

1. Khalaf WS, Mahmoud MRA, Elkhatib WF, Hashem HR, Soliman WE. Phenotypic characterization of NKT-like cells and evaluation of specifically related cytokines for the prediction of unexplained recurrent miscarriage. Heliyon. 2021 Nov 1;7(11):e08409.

2. Wang L, Deng Z, Yang J, Zhao Y, Zhou L, Diao L, et al. Epigenetic and transcriptomic characterization of maternal-fetal interface in patients with recurrent miscarriage via an integrated multi-omics approach. J Reprod Immunol. 2022 Dec 1;154:103754.

3. Stephenson M, Kutteh W. Evaluation and Management of Recurrent Early Pregnancy Loss. Clin Obstet Gynecol. 2007;50(1):132–45.

4. Popescu F, Jaslow CR, Kutteh WH. Recurrent pregnancy loss evaluation combined with 24-chromosome microarray of miscarriage tissue provides a probable or definite cause of pregnancy loss in over 90% of patients. Hum Reprod Oxf Engl. 2018 Apr 1;33(4):579–87.

5. Liu B, Wu H, Huang Q, Li M, Fu X. Phosphorylated STAT3 inhibited the proliferation and suppression of decidual Treg cells in unexplained recurrent spontaneous abortion. Int Immunopharmacol. 2020 May 1;82:106337.

6. Liang W, Chen M, Zheng D, Li J, Song M, Zhang W, et al. The Opening of ATP-Sensitive K+ Channels Protects H9c2 Cardiac Cells Against the High Glucose-Induced Injury and Inflammation by Inhibiting the ROS-TLR4-Necroptosis Pathway. Cell Physiol Biochem. 2017;1020–34.

7. Sun W, Wu X, Gao H, Yu J, Zhao W, Lu JJ, et al. Cytosolic calcium mediates RIP1/RIP3 complex-dependent necroptosis through JNK activation and mitochondrial ROS production in human colon cancer cells. Free Radic Biol Med. 2017;108:433.

8. Lu W, Sun J, Yoon JS, Zhang Y, Zheng L, Murphy E, et al. Mitochondrial Protein PGAM5 Regulates Mitophagic Protection against Cell Necroptosis. PloS One. 2016;11(1):e0147792.

9. Li CJ, Lin LT, Tsai HW, Wen ZH, Tsui KH. Phosphoglycerate mutase family member 5 maintains oocyte quality via mitochondrial dynamic rearrangement during aging. Aging Cell. 2022 Feb;21(2):e13546.

10. Lv Z, Xiong LL, Qin X, Zhang H, Luo X, Peng W, et al. Role of GRK2 in Trophoblast Necroptosis and Spiral Artery Remodeling: Implications for Preeclampsia Pathogenesis. Front Cell Dev Biol. 2021;9:694261.

11. Byun H, Kwon S, Wagner KU, Shin H, Lim HJ. Tumor susceptibility gene 101 is required for the maintenance of uterine epithelial cells during embryo implantation. Reprod Biol Endocrinol RBE. 2021 Jul 16;19(1):112.

12. Shui-Xing Y, Feng-Hua Z, Wei C, Gui-Mei J, Chong-Tao D, Gui-Qiu H, et al. Decidual Stromal Cell Necroptosis Contributes to Polyinosinic-Polycytidylic Acid-Triggered Abnormal Murine Pregnancy. Front Immunol. 2017;8:916.

13. Srinivasan S, Guha M, Kashina A, Avadhani NG. Mitochondrial dysfunction and mitochondrial dynamics-The cancer connection. Biochim Biophys Acta BBA - Bioenerg. 2017;1858(8):602–14.

14. Kai, Yasukawa, Hiroyuki, Oshiumi, Makoto, Takeda, et al. Mitofusin 2 inhibits mitochondrial antiviral signaling. Sci Signal [Internet]. 2009 [cited 2022 Dec 15]; Available from: http://stke.sciencemag.org/content/2/84/ra47

15. Mirjalili M, Mirzaei E, Vazin A. Pharmacological agents for the prevention of colistin-induced nephrotoxicity. Eur J Med Res. 2022 May 7;27(1):64.

16. Tuli HS, Aggarwal V, Kaur J, Aggarwal D, Parashar G, Parashar NC, et al. Baicalein: A metabolite with promising antineoplastic activity. Life Sci. 2020 Oct 15;259:118183.

17. Zhao WZ, Wang HT, Huang HJ, Lo YL, Lin AMY. Neuroprotective Effects of Baicalein on Acrolein-induced Neurotoxicity in the Nigrostriatal Dopaminergic System of Rat Brain. Mol Neurobiol. 2018 Jan;55(1):130–7.

18. Legarda D, Justus SJ, Ang RL, Rikhi N, Li W, Moran TM, et al. CYLD Proteolysis Protects Macrophages from TNF-Mediated Auto-necroptosis Induced by LPS and Licensed by Type I IFN. Cell Rep. 2016 Jun 14;15(11):2449–61.

19. Li Z, Scott MJ, Fan EK, Li Y, Liu J, Xiao G, et al. Tissue damage negatively regulates LPS-induced macrophage necroptosis. Cell Death Differ. 2016 Sep 1;23(9):1428–47.

20. Ihenacho UK, Meacham KA, Harwig MC, Widlansky ME, Hill RB. Mitochondrial Fission Protein 1: Emerging Roles in Organellar Form and Function in Health and Disease. Front Endocrinol. 2021;12:660095.

21. Weindel CG, Martinez EL, Zhao X, Mabry CJ, Bell SL, Vail KJ, et al. Mitochondrial ROS promotes susceptibility to infection via gasdermin D-mediated necroptosis. Cell. 2022 Aug 18;185(17):3214–3231.e23.

22. Dan Sang, Duan X, Yu X, Zang J, Liu L, Wu G. PGAM5 regulates DRP1-mediated mitochondrial fission/mitophagy flux in lipid overload-induced renal tubular epithelial cell necroptosis. Toxicol Lett. 2023 Jan 1;372:14–24.

23. Huang Z, Wang S, Yang Y, Lou J, Liu Z, Liu Z, et al. Mitochondrial dysfunction promotes the necroptosis of Purkinje cells in the cerebellum of acrylamide-exposed rats. Food Chem Toxicol. 2023 Jan 1;171:113522.

24. Faizan MI, Ahmad T. Altered mitochondrial calcium handling and cell death by necroptosis: An emerging paradigm. Mitochondrion. 2021 Mar 1;57:47–62.

25. Frank S, Gaume B, Bergmann-Leitner ES, Leitner WW, Robert EG, Catez F, et al. The role of dynamin-related protein 1, a mediator of mitochondrial fission, in apoptosis. Dev Cell. 2001 Oct;1(4):515–25.

26. Cheng M, Lin N, Dong D, Ma J, Su J, Sun L. PGAM5: A crucial role in mitochondrial dynamics and programmed cell death. Eur J Cell Biol. 2021 Jan 1;100(1):151144.

27. Cha J, Dey SK, Lim H (Jade). Chapter 38 - Embryo Implantation. In: Plant TM, Zeleznik AJ, editors. Knobil and Neill’s Physiology of Reproduction (Fourth Edition) [Internet]. San Diego: Academic Press; 2015 [cited 2022 Dec 28]. p. 1697–739. Available from: https://www.sciencedirect.com/science/article/pii/B9780123971753000387

28. Cha J, Sun X, Dey SK. Mechanisms of implantation: strategies for successful pregnancy. Nat Med. 2012 Dec;18(12):1754–67.

29. Grootjans S, Vanden Berghe T, Vandenabeele P. Initiation and execution mechanisms of necroptosis: an overview. Cell Death Differ. 2017 Jul;24(7):1184–95.

30. Fan X, Xu X, Wu X, Xia R, Gao F, Zhang Q, et al. The protective effect of DNA aptamer on osteonecrosis of the femoral head by alleviating TNF-α-mediated necroptosis via RIP1/RIP3/MLKL pathway. J Orthop Transl. 2022 Sep 1;36:44–51.

31. Zhu Y, Cui H, Xia Y, Hua G, Hiroyasu N. RIPK3-Mediated Necroptosis and Apoptosis Contributes to Renal Tubular Cell Progressive Loss and Chronic Kidney Disease Progression in Rats. Plos One. 2016;11(6):e0156729.

32. Robertson SA, Care AS, Moldenhauer LM. Regulatory T cells in embryo implantation and the immune response to pregnancy. J Clin Invest. 2018 Oct 1;128(10):4224–35.

33. Wang Z, Zibrila Abdoulaye I, Liu J, Li C. Trophoblast necroptosis in Preeclampsia: The role of fetus-derived exosomal microRNAs. Med Hypotheses. 2022 Oct 1;167:110949.

34. Yu H, Zhang Y, Liu M, Liao L, Wei X, Zhou R. SIRT3 deficiency affects the migration, invasion, tube formation and necroptosis of trophoblast and is implicated in the pathogenesis of preeclampsia. Placenta. 2022 Mar 24;120:1–9.

35. Guo J, Guo Q, Fang H, Lei L, Zhang T, Zhao J, et al. Cardioprotection against doxorubicin by metallothionein Is associated with preservation of mitochondrial biogenesis involving PGC-1α pathway. Eur J Pharmacol. 2014 Aug 15;737:117–24.

36. Chen W, Zhou Z, Li L, Zhong CQ, Zheng X, Wu X, et al. Diverse sequence determinants control human and mouse receptor interacting protein 3 (RIP3) and mixed lineage kinase domain-like (MLKL) interaction in necroptotic signaling. J Biol Chem. 2013 Jun 7;288(23):16247–61.

37. Wang H, Sun L, Su L, Rizo J, Liu L, Wang LF, et al. Mixed Lineage Kinase Domain-like Protein MLKL Causes Necrotic Membrane Disruption upon Phosphorylation by RIP3. Mol Cell. 2014 Apr 10;54(1):133–46.

38. Ryu MJ, Seo BJ, Choi YJ, Han MJ, Choi Y, Chung MK, et al. Mitochondrial and Metabolic Dynamics of Endometrial Stromal Cells During the Endometrial Cycle. Stem Cells Dev. 2020 Sep 24;

39. Hu J, Zhang Y, Jiang X, Zhang H, Gao Z, Li Y, et al. ROS-mediated activation and mitochondrial translocation of CaMKII contributes to Drp1-dependent mitochondrial fission and apoptosis in triple-negative breast cancer cells by isorhamnetin and chloroquine. J Exp Clin Cancer Res CR. 2019 May 28;38(1):225.

40. Meresman GF, Götte M, Laschke MW. Plants as source of new therapies for endometriosis: a review of preclinical and clinical studies. Hum Reprod Update. 2021 Feb 19;27(2):367–92.

41. Yu Z, Li Q, Wang Y, Li P. A potent protective effect of baicalein on liver injury by regulating mitochondria-related apoptosis. Apoptosis Int J Program Cell Death. 2020 Jun;25(5–6):412–25.

42. Deng X, Liu J, Liu L, Sun X, Huang J, Dong J. Drp1-mediated mitochondrial fission contributes to baicalein-induced apoptosis and autophagy in lung cancer via activation of AMPK signaling pathway. Int J Biol Sci. 2020;16(8):1403–16.

43. Jiang C, Zhang J, Xie H, Guan H, Li R, Chen C, et al. Baicalein suppresses lipopolysaccharide-induced acute lung injury by regulating Drp1-dependent mitochondrial fission of macrophages. Biomed Pharmacother Biomedecine Pharmacother. 2022 Jan;145:112408.

44. Pasparakis M, Vandenabeele P. Necroptosis and its role in inflammation. Nature. 2015 Jan 15;517(7534):311–20.

45. Zou H, Yin J, Zhang Z, Xiang H, Wang J, Zhu D, et al. Destruction in maternal-fetal interface of URSA patients via the increase of the HMGB1-RAGE/TLR2/TLR4-NF-κB signaling pathway. Life Sci. 2020 Jun 1;250:117543.

46. Zhang CS, Lin SC. AMPK Promotes Autophagy by Facilitating Mitochondrial Fission. Cell Metab. 2016 Mar 8;23(3):399–401.

47. Hosseinirad H, Paktinat S, Mohanazadeh Falahieh F, Mirani M, Karamian A, Karamian A, et al. Effect of 1,25(OH)2-vitamin D3 on decidualization of human endometrial stromal cells. Steroids. 2022 Apr 1;180:108978.

48. Wang K, Gao S, Wang J, Yu F, Ye C. Protective effects of chicoric acid on LPS-induced endometritis in mice via inhibiting ferroptosis by Nrf2/HO-1 signal axis. Int Immunopharmacol. 2022 Dec 1;113:109435.

49. Guo LT, Wang SQ, Su J, Xu LX, Ji ZY, Zhang RY, et al. Baicalin ameliorates neuroinflammation-induced depressive-like behavior through inhibition of toll-like receptor 4 expression via the PI3K/AKT/FoxO1 pathway. J Neuroinflammation. 2019 May 8;16(1):95.

